# Complete vertebrate mitogenomes reveal widespread gene duplications and repeats

**DOI:** 10.1101/2020.06.30.177956

**Authors:** Giulio Formenti, Arang Rhie, Jennifer Balacco, Bettina Haase, Jacquelyn Mountcastle, Olivier Fedrigo, Samara Brown, Marco Capodiferro, Farooq O. Al-Ajli, Roberto Ambrosini, Peter Houde, Sergey Koren, Karen Oliver, Michelle Smith, Jason Skelton, Emma Betteridge, Jale Dolucan, Craig Corton, Iliana Bista, James Torrance, Alan Tracey, Jonathan Wood, Marcela Uliano-Silva, Kerstin Howe, Shane McCarthy, Sylke Winkler, Woori Kwak, Jonas Korlach, Arkarachai Fungtammasan, Daniel Fordham, Vania Costa, Simon Mayes, Matteo Chiara, David S. Horner, Eugene Myers, Richard Durbin, Alessandro Achilli, Edward L. Braun, Adam M. Phillippy, Erich D. Jarvis, The Vertebrate Genomes Project Consortium

## Abstract

Modern sequencing technologies should make the assembly of the relatively small mitochondrial genomes an easy undertaking. However, few tools exist that address mitochondrial assembly directly. As part of the Vertebrate Genomes Project (VGP) we have developed mitoVGP, a fully automated pipeline for similarity-based identification of mitochondrial reads and *de novo* assembly of mitochondrial genomes that incorporates both long (>10 kbp, PacBio or Nanopore) and short (100-300 bp, Illumina) reads. Our pipeline led to successful complete mitogenome assemblies of 100 vertebrate species of the VGP. We have observed that tissue type and library size selection have considerable impact on mitogenome sequencing and assembly. Comparing our assemblies to purportedly complete reference mitogenomes based on short-read sequencing, we have identified errors, missing sequences, and incomplete genes in those references, particularly in repeat regions. Our assemblies have also identified novel gene region duplications, shedding new light on mitochondrial genome evolution and organization.

---

Mitochondria are found in the vast majority of eukaryotic cells^1^. The mitochondrial DNA (mtDNA) can be circular, as in animals, or linear, as in many plant species^2^. In animals, different cell types have varying numbers of mitochondria^3^, normally hundreds or thousands, with each mitochondrion usually harbouring 1-10 mtDNA copies^4^. In vertebrates, mtDNA varies from 14 to over 20 kbp in size, and albeit gene order can vary^5,6^, its gene content is highly conserved^2^. It usually contains 37 genes, encoding for 2 ribosomal RNAs (rRNAs), 13 proteins and 22 transfer RNAs (tRNAs). This “mitogenome” generally has short repetitive non-coding sequences, normally within a single control region (CR). However, relatively large repetitive regions, potentially heteroplasmic, have also been reported whose biological significance is still unclear^6,7^.

To date, mtDNA sequences have been generated for hundreds of thousands of specimens in many vertebrate species^8^. With some of its genes having been considered as the universal barcode of metazoa^9^, mtDNA is routinely employed at both the population and species levels in phylogeographic^10,11^, phylogenetic^12,13^, and paleogenomics studies^14,15^, among others^16,17^. The maternal inheritance pattern of mitochondria in vertebrates provides key complementary information to the nuclear DNA (nDNA), helping to reconstruct single maternal lineages without the confounding effects of recombination. The higher mutation rate of mtDNA compared to nDNA^16^, coupled with variable levels of conservation of mtDNA regions, can be used to delineate different types of phylogenetic relationships among species^18^. Key mtDNA molecular markers that have been used since the dawn of genetics include cytochrome b (*cob*), cytochrome c oxidase subunit I (*coi*), NADH dehydrogenase subunit 6 (*nad6*), and *16S* rRNA^19–21^. Thanks to its fast evolutionary rate, highly polymorphic nature, the non-coding CR has also been used in both present and ancient DNA studies^22^ to resolve the phylogenetic history of closely related species^23,24^.

At present, variation in the mtDNA sequence is usually assessed in two ways: 1) by target-enrichment and sequencing^25,26^; and 2) by *de novo* assembly from Whole-Genome Sequencing (WGS) with short reads^13^. In both scenarios, overlaps between sequences (Sanger or Next Generation Sequencing, NGS) have been used to assemble full-length mitogenomes. Despite the quantity and general high quality of mitogenomes reconstructions allowed by these methods, complex regions, particularly repeat regions and segmental duplications such as those sometimes present in the CR^27^, have been traditionally challenging to resolve^24,28^. Theoretically, whenever repeats longer than the reads are present, assemblies are limited within the boundaries of repetitive elements. Another important challenge for mitogenome assembly is posed by the nuclear DNA of mitochondrial origin (NUMT)^29,30^. NUMTs originate from the partial or complete transposition of the mtDNA into the nuclear genome, potentially leading to multiple copies of the mtDNA sequence scattered throughout it and independently evolving^31^. When probes, PCR primers, or *in silico* baits designed to match the mtDNA share sequence similarity with NUMTs, they can lead to off-target hybridization and amplification, resulting in the incorrect incorporation of NUMT sequence variation in mitogenome assemblies, ultimately impacting evolutionary analysis^29^.

In the last five years, novel WGS strategies based on single molecule, long-read sequencing technologies have been successful in improving the quality of the nuclear genome assemblies, particularly in repetitive regions^32^. Since single-molecule DNA sequencing can currently produce reads of at least 10-20 kbp in size^32^, a complete full-length representation of mtDNA genomes can theoretically be obtained in a single read, solving the overlap uncertainty issue of short-read NGS and Sanger-based approaches. However, since single molecule sequencing technologies have a relatively high error rate, assembly is still required to derive an accurate consensus sequence. Here, we describe a new mitochondrial genome assembly pipeline (mitoVGP) developed in the framework of the Vertebrate Genomes Project (VGP) (www.vertebrategenomesproject.org)^33,34^. The VGP aims to generate near complete and error-free reference genome assemblies representing all vertebrate species, and in its Phase 1 it is currently targeting one species for each vertebrate order. Our mitoVGP pipeline complements the nuclear assembly VGP pipeline^34^, which has become the current standard approach for all VGP species, by combining long reads for structural accuracy and short reads for base calling accuracy. We applied mitoVGP to determine the full mitogenome sequence of 100 vertebrate species, including 33 species for which a reference mtDNA sequence was not previously available. Compared to the published reference mitogenome assemblies, our assemblies filled gaps above 250 bp in size in 25% of cases, added missing repeats, genes or gene duplications, all of which lead to novel discoveries using these complete assemblies.

## Mitogenome assembly with mitoVGP

The VGP version 1 assembly pipeline developed for the nuclear genome uses Continuous Long Reads (CLR) and the Pacific Biosciences (PacBio) assembler FALCON to generate contigs^34,35^. When initially inspecting the contigs, we noted the absence of contigs representing the mitogenome in 75% of species. However, mitogenome sequences could be found in the raw reads, and we surmised they may have been filtered out due to size cutoffs or depth of coverage using nDNA assembly algorithms^36^, reducing the chances for the mitogenome to be represented in the final assembly. Moreover, when present in the final assembly, mtDNA contigs consist of long concatemers of the mtDNA sequence due to the circular nature of the mitogenome^37^. This issue motivated the development of mitoVGP, a novel bioinformatics pipeline specifically designed to obtain complete and error-free mitogenomes for all the species included in the VGP. MitoVGP uses bait-reads to fish out the mitogenome long reads from WGS data, assembles complete gapless mitogenome contigs, and polishes for base accuracy using short reads (**Extended Data Fig. 1;** workflow described in the **Methods**).

We evaluated the mitoVGP pipeline using paired PacBio long read and 10x Genomics WGS linked read datasets comprising 125 VGP vertebrate species belonging to 90 families and 59 orders (**Supplementary Table 1**). MitoVGP was able to generate mitogenome assemblies, with no gaps, in 100 cases (80%). The success rate was higher in some groups, such as mammals and amphibians, than in others, such as birds and fishes, albeit these lineage differences did not reach statistical significance (**Supplementary Table 2**; Fisher’s exact test, p = 0.12, N = 125). To test whether this difference was due to technical issues in the assembly pipeline or intrinsic properties of the raw data, we tested whether the assembly success was associated with the availability of long mtDNA reads. We found categorically that in all cases where long mtDNA reads were available the assembly was successful, while it failed in all cases where no long mtDNA reads were available (χ^2^ = 118.8, df = 1, p < 2.2 × 10^−16^, **Extended Data Fig. 2**).

A series of factors can theoretically affect the relative abundance of mtDNA reads, including taxonomic differences, tissue type, DNA extraction method, and library preparation protocols, particularly key steps such as size selection. We therefore fitted a linear model including all these factors (**Methods, Supplementary Table 4**). We found that the availability of mtDNA reads, and thus assembly success (**Supplementary Table 5**), varied significantly by *tissue type* (**Extended Data Fig. 3**), which explained the largest fraction of the total variance (25.3%). Mitogenome assembly using DNA derived from muscle was successful in 100% of the cases (N = 18). By contrast, blood (N = 34) and liver (N = 18) were used successfully only in 71% and 61% of the cases, respectively. The fraction of the variance in the number of mtDNA reads explained by *taxonomic group* (12.5%) was also significant, potentially as a consequence of real biological differences in mitochondria copy number among cells of different lineages. It should be noted that in our dataset *tissue type* and *taxonomic group* covaried, as seen in the preferential usage of some tissue types for specific taxonomic groups, such as nucleated blood for birds (**Supplementary Table 3**, χ^2^ = 302.8, simulated p = 9.999e-05). DNA extraction protocols (explained variance 1.2%) and library prep kits (1.1%) were not significantly associated with mtDNA read availability, but the library fragmentation approach (explained variance 6.9%) and size selection (11.7%) had significant effects. In particular, fragmentation with Megaruptor significantly preserved more mtDNA fragments than needle shearing (t = −2.767, p = 0.018) or no fragmentation (t = −3.387, p-value = 0.003), potentially because Megaruptor fragmentation can give a tight fragment size distribution whereas needles may over-shear the DNA and samples with low DNA integrity skipped fragmentation. While the relationship between the size selection cutoff used for the library preparation and assembly success is not monotonic, a significantly higher proportion of complete mitochondrial genomes were reconstructed from libraries with a size cutoff less than 20 kbp (χ^2^ = 16.6, simulated p = 3e-04, **Supplementary Table 6**). This is consistent with the mitogenome size usually being below 20 kbp, and library preparation protocols involving a higher cutoff would deplete the library of mtDNA. We also considered whether total sequence data impacted the availability of mtDNA reads; VGP datasets were all generated at approximately 60X CLR coverage of the estimated nuclear genome size. Coverage being equal, larger genomes (e.g. those of some amphibians > 4 Gbp) would generate considerably more raw data than smaller genomes (e.g. birds ∼ 1 Gbp). However, theoretically (see also **Supplementary Note 1**) and in practice (explained variance 0.3%) this did not translate into a substantial increase in the number of mtDNA reads.

Overall, the major role of read availability in assembly success, driven by the factors described above, supports the robustness and the unbiased nature of the mitoVGP pipeline in generating mitogenome assemblies in any genomic context.

## MitoVGP assemblies are more accurate and complete

The mitoVGP pipeline was developed under the assumption that a combination of long and short read datasets could provide a better representation of the mitogenome. K-mer-based QV estimates suggest a high base calling accuracy of mitoVGP assemblies (**Supplementary Table 1**, column AF), with 58 of the assemblies having no false mtDNA k-mers (i.e. false k-mers only found in the assembly and not in the high-coverage fraction of the raw data, see **Methods**), and the remaining with average QV 41.25 (approximately 1 base calling error per assembly). We confirmed the general high accuracy of mitoVGP assemblies generating orthogonal Nanopore long-read datasets on a subset of four VGP species using the same sample tissue. Overall, the two datasets for each species generated identical or near identical assemblies, both at the structural level and in base call accuracy (**Supplementary Table 7**).

We decided to benchmark the mitoVGP pipeline in the context of currently available mitogenome assembly tools using our VGP datasets. Unfortunately, the only alternative long-read organelle genome assembler to our knowledge, Organelle_PBA^38^, is no longer maintained nor functional (Aureliano Bombarely, personal communication). Therefore, we focussed on NOVOPlasty, a popular WGS short read mitogenome assembler^39^. Using NOVOPlasty on the short reads of all 125 VGP species, 70 assembled in a single contig, of which 61 were labelled as circular by NOVOPlasty, 48 assembled as multiple contigs, and the assembly failed in 7 cases (**Extended Data Fig. 4a**). NOVOPlasty did succeed in 24 cases where mitoVGP could not because of the absence of long mtDNA reads (15 assembled in multiple contigs, one in a single contig, 8 labelled as circular). When we compared the circular NOVOPlasty assemblies to their mitoVGP counterparts, we found the average level of identity in the alignable regions to be 99.913% (99.918% if IUPAC ambiguous base calls introduced by NOVOPlasty are not considered, **Fig. 1a**, green). There were 15 mitoVGP assemblies that were substantially larger than their NOVOPlasty counterparts, while all other assemblies had length differences < 5 bp (**Fig. 1b**, green). Interestingly, single-contig NOVOPlasty assemblies that were not labelled as circular (N = 8) were larger than their corresponding mitoVGP assemblies because they had failed to circularize, leaving large overlapping ends (**Fig. 1b**, yellow). Despite this, sequence identity levels were still remarkable: 99.978% (99.984% disregarding IUPAC bases in NOVOPlasty assemblies, **Fig. 1a**, yellow). In multiple-contig NOVOPlasty assemblies, the largest contig had an average identity of 99.764% (99.888% removing IUPAC bases, **Fig. 1a**, orange) with the respective mitoVGP assembly, and the average difference in length was 3,867 bp (**Fig. 1b**, orange), supporting the fragmented nature of some of these NOVOPlasty assemblies. Fragmented NOVOPlasty assemblies correspond to larger than average mitoVGP assemblies, suggesting that more complex mitogenomes were more problematic for NOVOPlasty (**Fig. 1b**, orange). The general high base-level identity observed is supportive of the high base calling accuracy of mitoVGP assemblies based on the combination of long and short reads. However, significant length divergence, particularly in the comparison with circular NOVOPlasty assemblies (one-sided paired samples Wilcoxon test, p-value = 0.0002), was due to the presence of few large indels. These large indels originate from an underrepresentation of repeats (**Fig. 1c**) and gene duplications (**Fig. 1d**) in the NOVOPlasty assemblies.

**Fig. 1.**
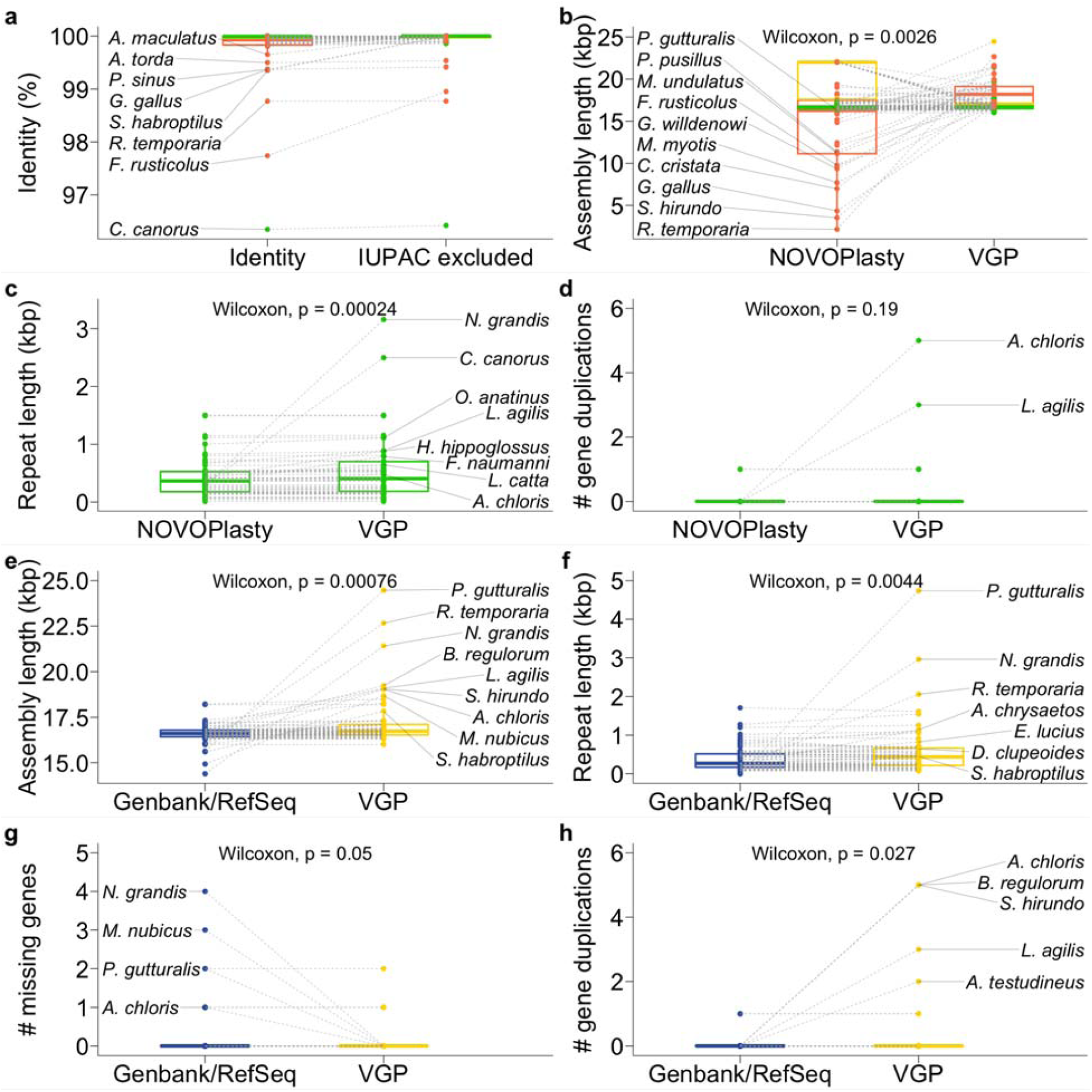
Paired comparisons of mitoVGP assemblies with NOVOPlasty and Genbank/RefSeq assemblies. NOVOPlasty assemblies are split into three categories: 1) circular (green); 2) single-contig (yellow); 3) multiple-contigs (orange). **a-d**, Comparisons between NOVOPlasty and mitoVGP assemblies for sequence identity (a, including and excluding IUPAC bases), assembly length (b), annotated repeat length (c, in circular NOVOPlasty assemblies and matched mitoVGP assemblies), and number of gene duplications (d, in circular NOVOPlasty assemblies and matched mitoVGP assemblies). **e-h**, Genbank/RefSeq comparisons of mitogenome assembly length (e), annotated repeat length (f), number of missing genes (g), and number of gene duplications (h). Statistical significance (one-sided paired samples Wilcoxon test) is reported above each plot. In the first plot outliers having identity <99.7% are labelled. Top10 outliers in the 20th and 80th percentiles are labelled in the other plots.

We compared the length of mitoVGP assemblies to their reciprocal Genbank/RefSeq counterparts, when available (**Extended Data Fig. 4b, Supplementary Table 1**, N = 66, RefSeq = 34, non-RefSeq = 32). MitoVGP assemblies were on average significantly longer (**Fig. 1e**). Similar to the NOVOPlasty comparison, this difference was mostly driven by mitoVGP assemblies having a significantly higher repeat content (**Fig. 1f**), particularly when they were longer than their Genbank/RefSeq counterparts (Spearman’s ρ correlation = 0.83, **Extended Data Fig. 5**), but only a marginally significant difference in GC content (**Extended Data Fig. 6**). MitoVGP assemblies also tended to have fewer missing genes (**Fig. 1h**), as well as a significantly higher representation of gene duplications (**Fig. 1g**).

## Novel duplications, repeats, and heteroplasmy

In the mitoVGP vs Genbank/RefSeq comparison, the top three outliers of mitogenome assembly length differences were found in birds (**Fig. 1e**): the yellow-throated sandgrouse (*Pterocles gutturalis*, reference generated using Illumina WGS^40^), the great potoo (*Nyctibius grandis*, long-range PCR and direct Sanger sequencing^41^), and the rifleman (*Acanthisitta chloris*, long-range PCR and direct Sanger sequencing^42^). These are indeed labelled as partial in Genbank, and lack most of the CR where repeats and gene duplications typically occur. In contrast, several other outliers have Genbank/RefSeq assemblies labelled as complete but are still shorter than their mitoVGP assembly counterparts. These include the sand lizard (*Lacerta agilis*), where in the mitoVGP assemblies (both Pacbio and Nanopore versions) we found a 1,977 bp-long repetitive duplication involving part of the origin of replication that comprises a tandem repeat (repeat unit = 36 bp, 199 bp-long), the terminal portion of *cob* gene, as well as *trnT* and *trnP* genes (**Fig. 2a**). Coverage profiles and repeat-spanning reads supported the presence and extent of the duplication. In particular, out of 145 Nanopore reads of average length 5,758 bp, 11 were above 16 kbp and fully spanned the repeat; similarly, of 36 PacBio reads of average length 8,153 bp, 6 were over 16 kbp-long and spanned the repeat. Due to the nature of PacBio libraries, the same molecule can potentially be read multiple times while the polymerase enzyme passes over the circular SMRTbell. We found at least 2 full-pass reads covering the same mtDNA molecule twice (forward and reverse strand), and the duplication was still present. This duplication was completely absent in our circular NOVOPlasty assembly (which is however 100% identical to the PacBio assembly for the rest of the sequence) as well as in the current RefSeq reference^43^ (**Fig. 2a**), strengthening the notion that short-read assemblies inevitably fail to represent gene duplications and repeats.

**Fig. 2.**
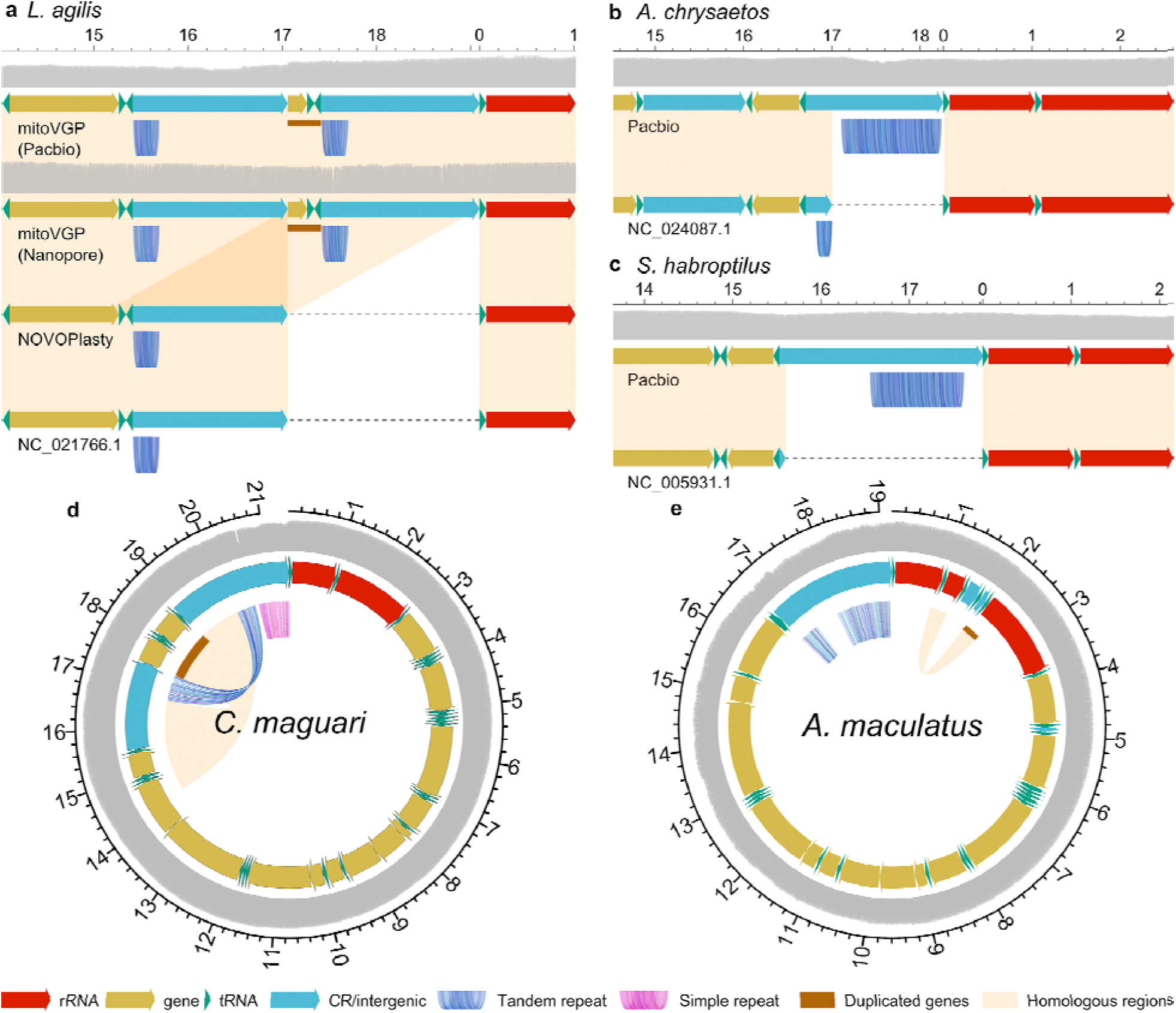
Duplications and repeats in mitoVGP assemblies. **a**, Comparison of mitoVGP, NOVOPlasty and RefSeq mitogenome assemblies for the sand lizard (*Lacerta agilis*). Duplicated genes missing from the reference and NOVOPlasty assemblies: *cob, trnT, trnP* (brown bar). Grey, read coverage: PacBio CLR 34x and Nanopore 46x mean coverage. **b**, Golden eagle (*Aquila chrysaetos*), where the current RefSeq sequence (top) lacks a large fraction of a tandem repeat in the CR and 10 bp from the start of the *trnF* gene (brown bar). Mean CLR coverage 170x. **c**, Kakapo (*Strigops habroptilus*), where the RefSeq sequence (top) lacks the entire CR. Mean coverage 99x. **d**, Maguari stork (*Ciconia maguari*). Duplicated genes: *cob, trnT, trnP, nad6* and *trnE* (brown bar). Mean CLR coverage 209x. **e**, Warty frogfish (*Antennarius maculatus*). Duplicated genes: *trnV* and *rrnL* (brown bar). Mean CLR coverage 21x. Th rRNA genes are colored in red, tRNA genes in green, other genes in yellow and the CR/intergenic region in blue. Homologous regions are highlighted in orange, tandem repeats in shades of blue, gaps as dashed lines and duplicated genes with brown bars. Long read coverage depth represented by the gray track. All labels in kbp. Coordinates relative to the PacBio mitoVGP assembly.

In the case of the golden eagle (*Aquila chrysaetos*), the 921 bp CR had a tandem repeat (repeat unit = 49 bp, ∼782 bp-long), which was essentially absent from the RefSeq reference (generated using Illumina WGS^44^) along with the first 10 bp encoding the *trnF* gene (**Fig. 2b**). Aside from the missing repeat, the remainder of the two sequences were 99.73% identical, the differences most likely attributable to individual variation. The high level of similarity is supportive of the overall quality of the mitoVGP assembly, which is also confirmed by its Q44.30 base call accuracy and 100% identity to the NOVOPlasty assembly in non-repetitive regions. In the Kakapo (*Strigops habroptilus*), an entire 2.3 kbp CR region, including a ∼925 bp-long repeat (repeat unit = 84 bp), was also missing from the RefSeq sequence (long-range PCR and direct Sanger sequencing^34,42^, **Fig. 2c**). In the case of the common tern (*Sterna hirundo*), the current RefSeq sequence (PCR and Sanger sequencing^45^) was missing a duplication in the CR involving the *cob, trnT, trnL2, nad6* and *trnE* genes, as well as a substantial fraction of a tandem repeat. In the case of the Indo-Pacific tarpon (*Megalops cyprinoides*), a ∼650 bp tandem repeat was represented in the RefSeq reference (long-range PCR and direct Sanger sequencing^46^) for two-thirds of its length, and the sequence downstream was completely absent, for ∼500 bp. The two sequences shared 99.89% identity, with mitoVGP assembly showing no base calling errors; our mitoVGP assembly was 100% identical to our NOVOPlasty assembly in the homologous regions, but the latter lacked 165 bp of the repeat, compatible with short reads falling short in covering the 650 bp repeat.

A total of 33 mitoVGP assemblies did not have a Genbank/RefSeq representative. Among these assemblies, 8 showed duplications and/or large repetitive elements. These include 5 birds belonging to 5 separate orders that have highly similar, but not identical patterns of duplicated genes in the CR (*cob, trnT, trnP, nad6* and *trnE*): the Anna’s hummingbird (*Calypte anna*)^34^, red-legged seriema (*Cariama cristata*), Swainson’s thrush (*Catharus ustulatus*), maguari stork (*Ciconia maguari*), whiskered treeswift (*Hemiprocne comata*) and European golden plover (*Pluvialis apricaria*). For example, in the maguari stork (*Ciconia maguari*, **Fig. 2d**) part of the CR itself is duplicated, and there is a simple repeat (CAA/CAAA, 842 bp-long) and two nearly identical tandem repeats (repeat unit = 70 bp, ∼523 bp-long and ∼719 bp-long respectively), leading to a 21.4 kbp-long mitogenome (assembly QV 41.41). Another example of newly identified gene duplications is represented by the warty frogfish (*Antennarius maculatus*, **Fig. 2e**). Here *trnV* and *rrnL* genes are partially duplicated while the CR shows two distinct repetitive elements, a short tandem repeat (repeat unit = 14 bp, ATA<u>AC</u>ATACATTAT / ATA<u>GT</u>ATACATTAT, 1,330 bp-long) and a longer tandem repeat (repeat unit = 54 bp, ∼479 bp-long), leading to a mitogenome size of 19.2 kbp with no base calling errors.

Overall, over 50% of mitoVGP assemblies (52/100) presented one or more repeats (N = 45) and/or duplications (N = 18) (**Fig. 3a, Supplementary Table 1**, columns AJ-AQ). Repeats, mostly in the CR, were detected in a higher proportion of reptiles (100%, 4/4, average length = 267 bp, sd = 178 bp), followed by amphibians (50%, 3/6; average length = 904; sd = 484 bp), mammals (46.2%, 12/26; average length 308 bp, sd = 129 bp), birds (44.4%, 12/27; average length = 1,181 bp, sd = 830 bp), and least in fish (37.5%, 12/32; average length = 549 bp; sd = 352). Duplications were in the highest proportion in birds (37%, 10/27), followed by reptiles (25%, 1/4), fish (12.5%, 4/32), and least in mammals (11.5%, 3/26). Repeats in the CR were often present when a duplication was found. According to the phylogeny and given the high variability among species, some repeats and duplications within and across vertebrate lineages can be interpreted to have diverged from a common ancestor or have independently converged (**Fig. 3a**). For example, since most birds sequenced to date have repeats in the CR, we can infer that it was likely the ancestral state, but then lost in some lineages and widely diverged in others. For duplications, similar to what has been claimed for Passeriformes^27^, duplications in or near the control region were present in several monophyletic clades that represent basal branches, spanning the Caprimulgiformes (e.g. hummingbird, *Calypte anna*) to Charadriiformes (e.g. razorbill, *Alca torda*), suggesting that this duplication was ancestral among them, and lost in close relatives (e.g. great potoo, *Nictus grandis*; **Fig. 3a**). These and other hypotheses on the phylogenetic history of mitochondrial repeats and duplications will be more quantitatively resolved once the remaining ordinal level genomes of the VGP Phase 1 are completed.

**Fig. 3.**
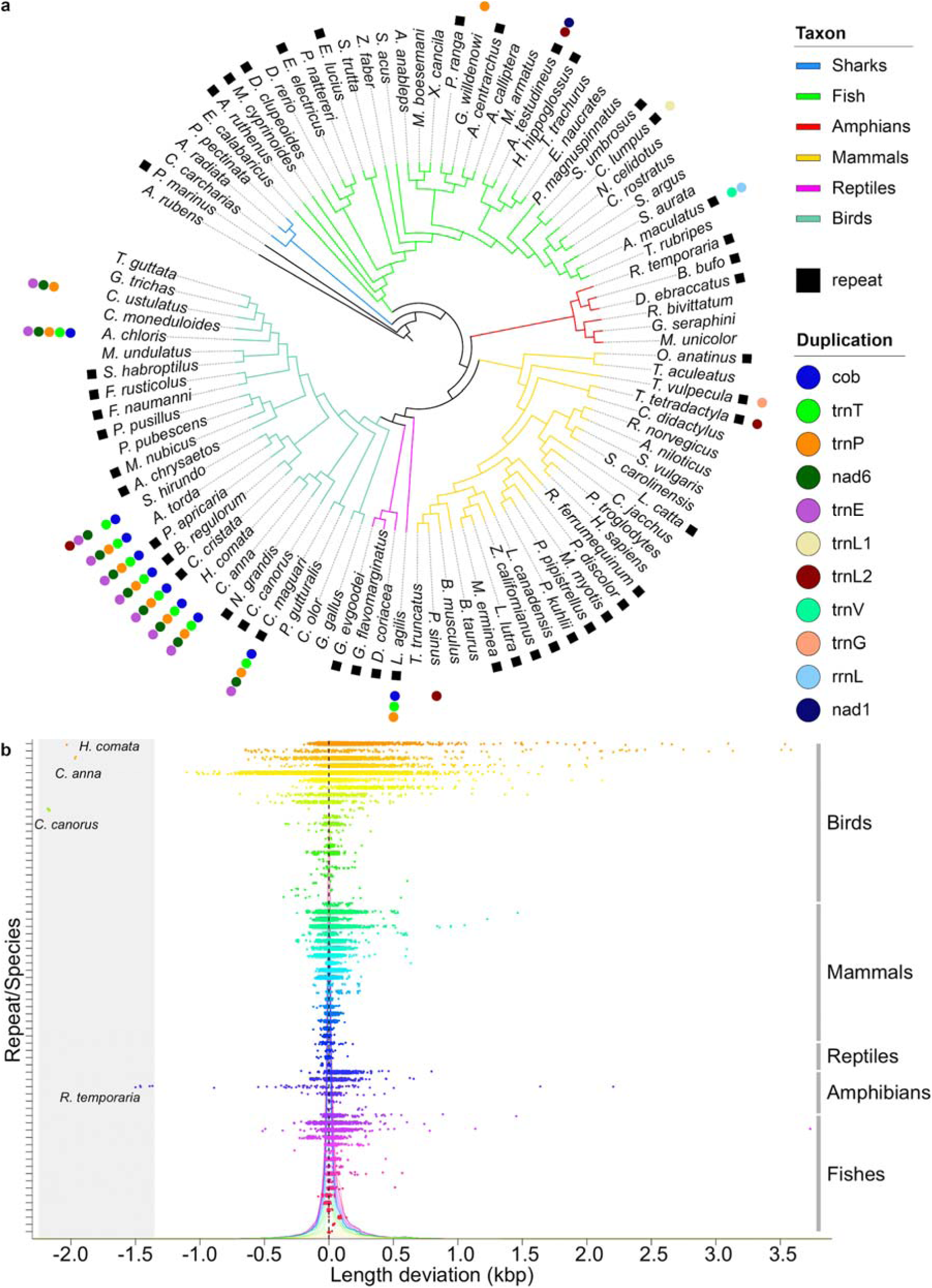
Duplications and repeats across the phylogeny and length heteroplasmy deviation in repetitive elements. **a**, The presence of mitochondrial repeats and duplications are mapped onto the tree for each species. The repeats are most often in the CR. Since the tree topology of phylogenies based on mtDNA is often inaccurate, this tree topology is based on relationships determined from current genome-scale phylogenies in the literature^49–52^. **b**, The length deviation from the reference is reported for each read spanning the repeat region. No deviation from the assembled VGP reference is marked by the dashed line. Colors correspond to different repetitive elements. Individual density distributions are shown in the background. The gray shaded area highlights four species that had reads that lack gene duplications when these are present in the mitoVGP assembly, suggesting possible heteroplasmy.

Long-read platforms offer the additional advantage of single molecule-resolution without amplification bias. To make sure that our assemblies represented the most frequent allele in the presence of heteroplasmy and assess the degree of heteroplasmy, we measured individual read length variation in repetitive elements and duplications. The analysis revealed that the read length difference from the VGP reference allele is centered around zero (**Fig. 3b**), with generally minor deviations largely explained by the high indel error rate of PacBio reads. Positive deviations were significantly favored (t = 29.631, df = 38374, p-value < 2.2e-16), compatible with the higher number of insertions over deletions in PacBio indel errors^47^. In some cases, a number of extra or missing copies of repetitive elements were present in individual reads (e.g. **Extended Data Fig. 7**), and the standard deviation linearly correlated with the size of the repeat element (Spearman’s ρ correlation = 0.79, p-value < 8.1e-16). Again, positive deviations were favored, suggesting an insertion bias in larger repeats. Interestingly, the duplications involving multiple genes in the CR also showed evidence of limited heteroplasmy in a few cases (**Fig. 3b**, gray area). Birds tended to have greater length deviations than the rest of vertebrates, indicating that they may have higher levels of heteroplasmy. The presence of repeats or duplications was not associated with tissue type (repeats: Fisher’s exact test, p-value = 0.11, N = 125; duplications: Fisher’s exact test, p-value = 0.16, N = 125), suggesting that these are not transient, tissue-specific events.

Of note, the advantage of single-molecule resolution is especially evident with the recently developed PacBio HiFi (High-Fidelity) technology, which generates ∼10-20 kb long reads having lower error profiles close to Illumina short reads^48^. To evaluate the accuracy attainable with these datasets, we generated HiFi reads for the human trio VGP dataset and mapped them to our reference mitogenome. The alignments showed uniform coverage across the assembly, easily allowing the identification of SNPs and indels at single molecule resolution and therefore greatly expanding the potential for heteroplasmy detection (**Extended Data Fig. 8**).

## Length distribution of vertebrate mitogenomes

Having ascertained that several of our assemblies are likely more complete than the respective existing references, we compared the mtDNA sequence length distribution of mitoVGP assemblies with the overall distribution of RefSeq representative mitogenomes of vertebrates labelled as complete. A final RefSeq dataset containing 3,567 sequences, excluding our VGP submissions (**Supplementary Table 8**), was randomly resampled to ensure the same within-order representation of sequences as the VGP dataset (1,000 replicates). The two datasets consistently diverged for lengths >18 kbp, with longer sequences represented in the mitoVGP dataset (**Fig. 4**). This suggests that the underrepresentation of repeats and duplications observed in our sample is likely to affect many of the existing reference assemblies deemed complete. It also highlights the multimodal distribution of mitogenome length in vertebrates, with mitoVGP assemblies revealing at least a secondary peak due to the presence of repeats and duplications.

**Fig. 4.**
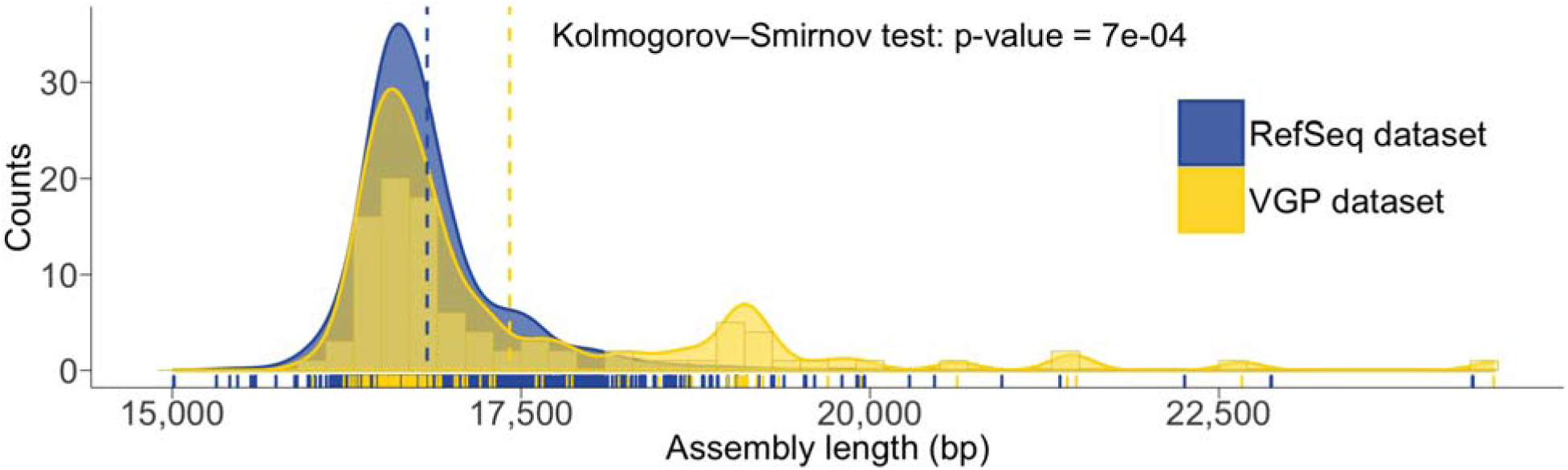
Distribution of vertebrate mtDNA sequence lengths in the VGP and RefSeq datasets. Length distribution histogram in the VGP dataset (yellow bars), with its density distribution (yellow area) and the RefSeq dataset density distribution (blue area). The RefSeq dataset was randomly resampled to ensure the same within-order representation of sequences as the VGP dataset (1,000 replicates). The respective means are highlighted by the dashed lines. Individual data points are shown at the bottom.

## Recommendations and evolutionary interpretations

We have demonstrated that mitoVGP, a new mitogenome assembler combining long and short reads, can successfully assemble high-quality mitogenomes in a variety of datasets. The number, quality and variety of complete mitogenome assemblies and datasets presented here allows a more accurate comparison of sequence data and assembly strategies than ever before. Our results suggest that when targeting the mitogenome, a careful design should be used to decide the sequencing technology, with further attention paid to the evaluation of assembly results. Tissue types with abundant mtDNA and libraries that avoid too stringent size selection should be preferred to ensure the presence of mtDNA reads in WGS experiments with long reads. Possibly due to their shallow size selection, current Nanopore library preparation protocols favor the availability of mtDNA reads over current PacBio CLR library protocols (see **Methods**). Alternatively, if stringent size selection is avoided, given their length and base accuracy, HiFi PacBio reads are an excellent candidate for future mitogenome studies on health and disease of vertebrate species, providing incontrovertible single molecule assessment.

The Genbank nucleotide database, one of the most complete DNA sequence archives, contains thousands of animal mtDNA sequences. A search in the animal subset (June 2020, keywords “mitochondrion AND complete”), yielded over 100,000 mitochondrial sequences, of which 75,565 were vertebrates. When the sequence is reported as *complete* in the metadata, we suggest it should imply that the full mtDNA sequence, circular in the case of vertebrate mitogenomes, has been assembled with no gaps. However, when compared with our long-read mitogenome assemblies, a proportion (at least 15%) of short-read assemblies publicly available in Genbank/RefSeq repositories and labelled as complete, are missing repeats and gene duplications. Unfortunately, these assemblies usually lack the publicly available supporting raw sequence data, making it difficult, or at times impossible, to evaluate their quality. Despite this limitation, we have shown that the discrepancy between mitoVGP and previous submitted Genbank/RefSeq assemblies is due to the use of long reads that span repetitive elements and duplicated genes. The presence of repeats and duplications in over half of the species herein assembled indicates that their occurrence is a principle of mitochondrial structure rather than an exception. Given the relatively high frequency of these elements, even in the “simple” case of vertebrate mitogenomes, the completeness of many currently available reference sequences can be further improved. Therefore, caution should be exercised before claiming complete assembly of a mitogenome from short reads alone.

## Methods

### VGP data generation

PacBio and 10x Genomics datasets for all VGP species were generated following protocols detailed in our companion paper on the nuclear assembly pipeline^34^ and in Mountcastle et al. (in preparation). The final dataset (N = 125) includes one invertebrate (the common starfish, *Asterias rubens*) and two individuals for the zebra finch (*Taeniopygia guttata*, one male and one female). A summary of the approaches employed for the samples analyzed in this work is provided in **Supplementary Table 1**. Briefly, total genomic DNA (gDNA) was obtained using a variety of state-of-the-art approaches for High Molecular Weight (HMW) DNA extraction available mostly at three different sequencing facilities of the contributing to the VGP (https://vertebrategenomesproject.org/): The Rockefeller University Vertebrate Genome Laboratory in New York, USA; the Wellcome Trust Sanger Institute in Hinxton, UK; and the Max Planck Institute in Dresden, Germany. This includes the Bionano plug protocol for soft tissue (Cat. No. 80002) and nucleated blood (Cat. No. 80004), MagAttract HMW DNA Kit for blood and tissue (Cat. No. 67563) and Phenol-Chloroform extraction. Library preparation followed standard protocols as suggested by the datasheets. In several cases, the DNA was fragmented using the Megaruptor at various fragment sizes between 15 and 75 kbp. In other cases, the DNA was fragmented by needle shearing. Importantly, libraries were usually size-selected to enrich for HMW fragments, and the range of size selection varied widely between 7 to 40 kbp. Both PacBio CLR and 10x Genomics linked reads were generated for all species in the VGP dataset, except the common starfish (*Asteria rubens*) and the chimp (*Pan troglodytes*) for which 10x was replaced with standard Illumina library preparation and publicly available data (SRX243527), respectively. For the human trio, we generated ∼10 kbp CCS libraries for all samples.

### Nanopore data generation

For the Nanopore datasets, total gDNA was obtained from tissue using Genomic-tip 100/G (Qiagen) for the spotty (*Notolabrus celidotus*), thorny skate (*Amblyraja radiata*), and hourglass treefrog (*Dendropsophus ebraccatus*), following the manufacturer’s protocol. For the sand lizard blood, total gDNA was extracted using Nanobind CBB Big DNA kit (Circulomics) as described by the manufacturer. All extracted gDNA underwent size selection using the Short Read Eliminator kit (Circulomics) to deplete fragments <10kb. The resulting material was then prepared for sequencing using the Ligation Sequencing Kit (SQK-LSK109, Oxford Nanopore Technologies Ltd) and sequenced using R9.4.1 flowcells (FLO-PRO002) on the PromethION device (Oxford Nanopore Technologies Ltd). Flowcell washes and library re-loads were performed when required. Interestingly, at matched coverage most Nanopore datasets contained a larger amount of mtDNA reads compared to PacBio (average fold change 3.9, **Supplementary Table 7**). The sole exception was the thorny skate, where the two datasets are comparable (1.6 fold more reads in the PacBio dataset). The PacBio dataset for the thorny skate was one of the very first VGP datasets produced, and at that time size selection was not as stringent.

### Mitogenome assembly pipeline

The mitoVGP pipeline was designed to simultaneously take advantage of the availability of long reads (especially PacBio) and short reads (usually 10x linked reads) data from the same individual. This condition is met for all genomes sequenced under the VGP nuclear genome pipeline, as well as for Nanopore datasets. The pipeline is fully automated, and it is composed of a series of single-node, fully parallelized, Bash scripts designed to run in a Linux environment. The amount of resources required is minimal, and will only affect speed. Command line code to reproduce our results for each assembly using mitoVGP is provided in **Supplementary Table 1**.

Similar to other methods available to assemble organelle genomes^38,39,53^, mitoVGP general workflow starts by selecting putative mitochondrial reads from a long-read WGS dataset based on their similarity with an existing reference of the same or other species, even distantly related ones. In mitoVGP 2.2, this is achieved using pbmm2 v1.0.0 (https://github.com/PacificBiosciences/pbmm2), the official PacBio implementation of Minimap2 (https://github.com/lh3/minimap2)^54^ for PacBio long reads, and using directly Minimap2 v2.17 for Nanopore reads. MitoVGP currently supports a variety of different PacBio chemistries. For RSII chemistries the aligner blasr v5.3.3 was employed via the -m option (https://github.com/PacificBiosciences/blasr). Reads were individually aligned to a mtDNA reference sequence using default parameters and allowing for unique alignments. The reference can be of the same species, or that of a closely-to-distantly related species, since even with default parameters pbmm2/Minimap2/blasr search similarity cut-offs are relatively loose, to account for the high error rate of noisy long reads^54^. We have experimentally determined in one VGP dataset (Anna’s hummingbird) that, when mapping with pbmm2 and *C. elegans* mitogenome indel/substitution rates^55^, an edit distance between the reference and the sample as large as 20% will decrease the number of available reads by 41% (total Gbp decrease 25%), and therefore having no substantial impact in long read availability for assembly in most cases. For comparison, mouse (*Mus musculus*, NC_005089.1) and zebrafish (*Danio rerio*, NC_002333.2) edit distance is 33.1%. The use of even more distant reference sequences should be possible using lower stringency in the mapper. Moreover, such references do not have to be complete, since even short matches on a fragmented reference will allow fishing out long reads, potentially spanning the gaps in the reference.

The next step of the pipeline involves the *de novo* genome assembly of the long reads extracted from the WGS dataset using the long read assembler Canu v1.8 (https://github.com/marbl/canu)^36^. After the assembly, since the Canu output may contain more than one contig, we used BLAST^56^ to identify and filter out contigs originating from lower quality mtDNA reads or failed overlaps as well as from the inclusion of nDNA reads. The sequence of the putative mitocontig was then polished using Arrow (https://github.com/PacificBiosciences/GenomicConsensus) in the case of PacBio datasets, or one round of Racon (https://github.com/isovic/racon) and one of Medaka (https://github.com/nanoporetech/medaka) in the case of Nanopore datasets. The sequence was further refined with a round of polishing using short read data, where the WGS short reads were mapped with Bowtie2 v2.3.4.1^57^ to extract putative mtDNA reads, variant calling on the alignments performed with Freebayes v1.0.2^58^, and consensus generated with Bcftools v1.9 using parameters optimal for a haploid genome and to left-align and normalize indels.

Because Canu was developed to assemble large linear nuclear chromosomes rather than short circular genomes, it leaves large overlaps at the sequence ends^36^. In a few cases, particularly when PacBio reads were missing adapter sequences resulting in subreads having multiple copies of the same DNA sequence, these overlaps can become exceptionally long, including the mitochondrial sequence read through multiple times. A custom script was developed to remove the overlaps (https://github.com/gf777/mitoVGP/blob/master/scripts/trimmer). Contrary to other methods that are based on the sequence of the overlaps alone (e.g. Circlator^37^ https://sanger-pathogens.github.io/circlator/), the script is based on the deconvolution of the repetitive elements using MUMmer matches (https://mummer4.github.io/)^59^, short-read mapping to the sequence using Bowtie2^57^ and coverage-based definition of the reliable ends. A final round of short-read polishing was performed using the same approach as previously described. Although we used 10x Genomics linked short-reads, mitoVGP accepts any short-read dataset in FASTQ format. Finally, tRNAs were identified using tRNA-scan-SE^60^, and the sequence was automatically oriented to start with the conventional tRNA Phenyl-Alanine (trnF).

### mitoVGP parameters

For 34 species, default mitoVGP parameters were adjusted as detailed in **Supplementary Table 1**. Relevant parameters include a query coverage cutoff to filter spurious BLAST^56^ matches during the identification of the putative mitochondrial contig (-p option), the maximum read length cutoff applied to long reads extracted with the aligner in the first step (-f option), Canu defaults (which can be adjusted directly in mitoVGP using the -o option) and the MUMmer cutoff for extending matches (-s option). The query coverage cutoff is important to identify and remove reads that only share a very loose similarity with the reference, such as repeats or small common motifs. Very short spurious matches can be filtered with a small value (e.g. -p 5), but if the reference is considered complete and closely related, query coverage can safely be increased (e.g. -p 70) as it is generally robust to the low accuracy of long reads. Filtering out reads that are significantly longer than the expected mitogenome assembly size is a complementary strategy that in some cases can considerably improve the quality of the assembly, since these reads are most likely NUMTs or nuclear repeats. Canu defaults can be changed for a variety of reasons, such as different error rates due to chemistry or quality of the dataset. When only a few mtDNA reads are available, the option *stopOnLowCoverage* needs to be tweaked (e.g. -o “stopOnLowCoverage=9”). The MUMmer cutoff should normally be adjusted only in a few cases, for instance in the presence of very large repeats. Nanopore-based assemblies were generated using the same pipeline. For the spotty and the hourglass treefrog mitoVGP was run with default parameters. For the sand lizard the -f option was adjusted to 25,000 and -p was adjusted to 5. For the thorny skate the -f option was adjusted to 18,000.

### Statistical analyses

Fisher’s exact tests and χ^2^ tests were performed with the respective primitive R functions using default parameters. Fisher’s exact test was used when the counts in each category were not sufficient for a reliable χ^2^ test. In χ^2^ tests, simulated p-values were generated using 10,000 replicates.

The set of reliable mtDNA reads used for statistical analyses was defined by the reads that covered the reference by at least 70% of their length using BLAST^56^ matches using the reference as query and the reads as reference (formula: reference length/read length*query cover). For three species with phylogenetically distant reference sequences, i.e. largescale four-eyed fish (*Anableps anableps*), warty frogfish (*Antennarius maculatus*) and blunt-snouted clingfish (*Gouania willdenowi*), the mitoVGP assembly was used to extract the reads to avoid underestimating read counts.

The linear model was implemented in R using the primitive R function *lm*:

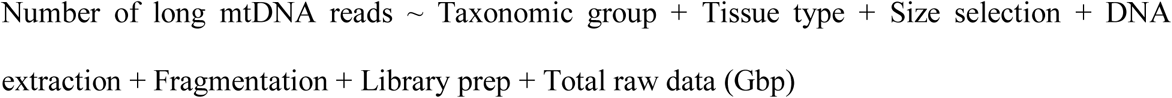

The fraction of the variance explained by each predictor was computed as Sum Sq/Total Sum Sq*100. For size selection, when multiple libraries with different cutoffs were generated, the minimum cutoff was considered in the statistical analyses. All observations having a factor represented 3 times or less were excluded (N=94). Post-hoc tests were performed with the *glht* function in the multcomp R library. We carefully checked model assumptions and further checked statistical significance of all terms with a permutation approach using the *lmp* function in the lmPerm R library. Results were always consistent, so for simplicity we report only the results of the parametric test.

### Measure of QV

QV of the mitogenome assemblies was assessed using a recently developed k-mer method called Merqury (https://github.com/marbl/merqury)^61^. The tool was run with default parameters and k-mer size 31, but given the approximate 50-100x coverage of VGP datasets, k-mers with frequency <100 were removed, to limit QV overestimation due to the inclusion of nDNA k-mers. 15 datasets were excluded from the analysis since mitogenome short read coverage <250x (**Supplementary Table 1**, column AE) overlapped substantially with the nDNA k-mers distribution and was similarly affected by the cutoff, preventing a reliable QV estimate.

### Benchmarking with existing tools

We run NOVOPlasty^39^ with default parameters, using the same reference sequence employed for mitoVGP as bait, and trimming 10x datasets using proc10xG with the -a option (https://github.com/ucdavis-bioinformatics/proc10xG). Following NOVOPlasty authors’ suggestion and in order for the results to be comparable to mitoVGP assemblies, the entire short read set was employed. Despite the fact that both mitoVGP and NOVOPlasty assemblies run on machines with 32 cores and 383 GiB of memory, in the case of NOVOPlasty the large size of VGP datasets often led to memory issues as all the reads have to be simultaneously loaded into a hash table. Assemblies that initially failed with NOVOPlasty were rerun on two fat nodes, each with 64 cores and 1.534 and 3.096 TB of memory respectively, narrowing down to four the number of failed assemblies due to reproducible internal errors (European Toad, Eurasian blackcap, cow, Canada lynx). Among the successful assemblies, one assembly (Anna’s hummingbird) was made of multiple contigs < 3 kbp and < 90% identity with the reference, one assembly (stoat) was made of multiple contigs with no identity with the reference, and one assembly (common pipistrelle) was made of a single 253 bp contig. These results were likely due to the inability of NOVOPlasty to identify an appropriate seed, and these assemblies were excluded from downstream analyses.

### Annotation

In all datasets, genes were annotated using MITOS v2.0.6 (https://gitlab.com/Bernt/MITOS), with default parameters. For the VGP and Genbank/RefSeq comparison, since MITOS may sometimes generate two contiguous annotations out of a single gene, the annotations were manually curated to identify real missing genes and gene duplications. Simple and tandem repeats were annotated using WindowMasker v1.0.0, with default parameters. K-mers were counted combining both the mitoVGP assembly and its Genbank/RefSeq or NOVOPlasty counterpart, and the difference in repeat representation between the two datasets was measured as the difference in the number of bases annotated as repetitive for each species.

### Coverage tracks

Reads were remapped on the original assemblies and filtered for identity to the reference >70% and length above ∼ the repeat length + 2,000 bp to ensure that they could anchor on both sides of the repetitive elements, avoiding contamination by nuclear repeats. For circos plots, reads were mapped to a 2-copy concatemer of the reference sequence two allow accurate read mapping at the edges.

### Phylogenetic tree

For the consensus tree, Timetree (http://www.timetree.org/)^62^ was initially used to generate a topological backbone of all species with a mitoVGP assembly. The tree topology was edited using TreeGraph2^63^ to reflect the current hypotheses on the phylogeny of the major vertebrates clades^49–52^.

### Measure of heteroplasmy

The coordinates of repeats and duplications were identified by manual inspection of BLAST^56^ off-diagonal matches with no repeat masking. The deviation from the reference was assessed by mapping the full WGS read set to the reference with pbmm2 v1.0.0. Confident mtDNA reads were identified in the same way used to define the read set for the linear model, the reads fully overlapping the repeat were filtered using bedtools^64^ and hard-clipped with jvarkit^65^.

### RefSeq sequence analysis

In order to collect RefSeq mtDNA sequences from Genbank, on March, 30th 2020 we conducted a custom search using the following keywords *“mitochondrion AND srcdb_refseq[PROP] AND complete [TITLE] NOT complete cds[TITLE] NOT isolate NOT voucher_03302020”*. To assign taxonomy and generate random samples belonging to the same orders of the VGP datasets we used the R package t*axonomizr*. Within-order resampling was conducted with 1,000 replicates.

### Graphical representations

For pairwise comparisons, we used R packages *ggplot2, gtable* and *ggrepel*. To represent gene duplications in the VGP and Genbank/RefSeq comparison, we used the R package *shape*. For Circos plots, we used the R package *circlize*. The phylogenetic tree was annotated with iTOL (https://itol.embl.de/)^66^. For density plots, we used the R package *ggplot2*. For the correlation, we used R packages *ggplot2* and *ggpubr*. Figures of read alignments were generated using IGV^67^.

## Data availability

The mitogenome assemblies are made available in the VGP GenomeArk (https://vgp.github.io/genomeark/) and under the NCBI/EBI BioProjects that will be listed in **Supplementary Table 1** upon publication. The nuclear genome assemblies of these species are current under the G10K embargo policy https://genome10k.soe.ucsc.edu/data-use-policies/.

## Code availability

MitoVGP is available under BSD 3-Clause License through the VGP Github portal (https://github.com/VGP/vgp-assembly) along with a ready-to-use conda environment, fully commented code, complete instructions and examples on how to run it. The code to reproduce the analyses and the plots is available at https://github.com/gf777/mitoVGP.

## Acknowledgments

We thank the contributors of the VGP on the first 125 species for letting us use data for generating mitochondrial genome assemblies; in particular they are Alexander N. G. Kirschel, Andrew Digby, Andrew Veale, Anne Bronikowski, Bob Murphy, Bruce Robertson, Clare Baker, Camila Mazzoni, Christopher Balakrishnan, Chul Lee, Daniel Mead, Emma Teeling, Erez Lieberman Aiden, Erica Todd, Evan Eichler, Gavin J.P. Naylor, Guojie Zhang, Jeramiah Smith, Jochen Wolf, Justin Touchon, Kira Delmore, Kjetill Jakobsen, Lisa Komoroske, Mark Wilkinson, Martin Genner, Martin Pšenička, Matthew Fuxjager, Mike Stratton, Miriam Liedvogel, Neil Gemmell, Piotr Minias, Peter O. Dunn, Peter Sudmant, Phil Morin, Qasim Ayub, Robert Kraus, Sonja Vernes, Steve Smith, Tanya Lama, Taylor Edwards, Tim Smith, Tom Gilbert, Tomas Marques-Bonet, Tony Einfeldt, Byrappa Venkatesh, Warren Johnson, Wes Warren, and Yury Bukhman. We are grateful to the Ngā Papatipu Rūnanga o Murihiku and the Ngāi Tahu for their support in generating the kakapo datasets. We also thank Aureliano Bombarely for his support in testing Organelle_PBA on VGP datasets. A. R., S. K., and A. M. P. were supported by the Intramural Research Program of the National Human Genome Research Institute, National Institutes of Health. A. R. was also supported by the Korea Health Technology R&D Project through KHIDI, funded by the Ministry of Health & Welfare, Republic of Korea (HI17C2098). F. O. A. was supported by Al-Gannas Qatari Society and The Cultural Village Foundation-Katara, Doha, State of Qatar and Monash University Malaysia. G. F. and E. D. J were supported by Rockefeller University start-up funds and the Howard Hughes Medical Institute.

## Author contributions

G. F. built the mitoVGP pipeline with support from A. R., D. H., M. C., S. K., S. M., A. F., A. M. P. and the mitoVGP assembly working group. G. F. performed the bioinformatic and statistical analyses with support from F. O. A., R. B., E. B, K. H., A. T., J. W., M. C., P. H., A. A., J. K., J. T. and the mitoVGP assembly working group. PacBio and 10x data were generated by O. F., B. H., J. M., I. B, J. B., S. W, K. O., J. S., E. B., J. D., M. S. and C. C. Nanopore datasets were provided by V.C., S. M. and D. F. W. K. and J. T. helped with metadata collection and submission of the assemblies to the public archives. G. F. and E. D. J. conceived the study and drafted the manuscript. All authors contributed to the final manuscript and approved it.

## Materials & Correspondence

Please address any request to Dr. Erich Jarvis and Dr. Giulio Formenti, The Rockefeller University, New York, NY, USA.

## Extended data figures

**Extended Data Fig. 1.**
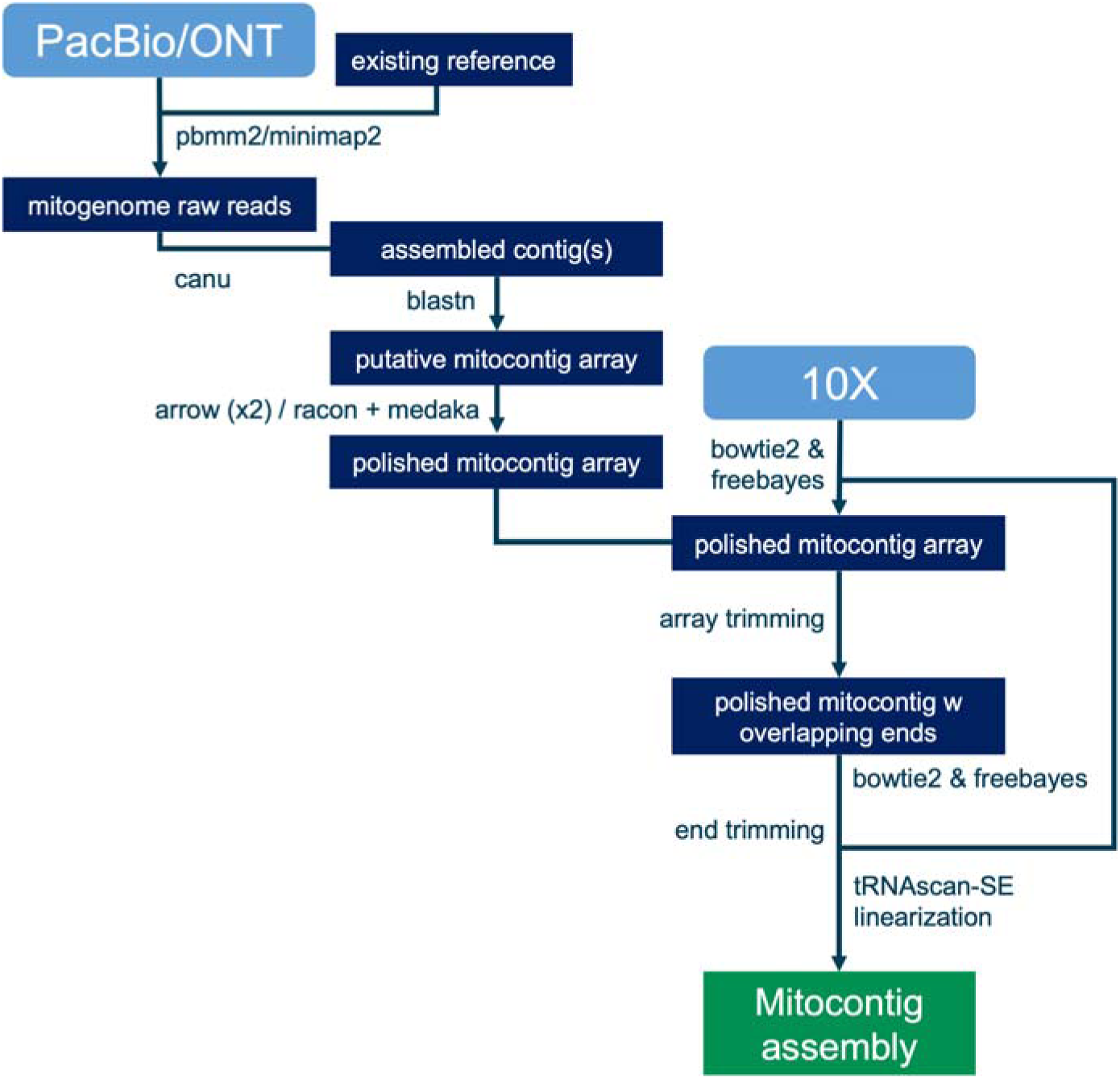
Outline of the mitoVGP assembly pipeline. Raw read data are represented by light blue rectangles. The outputs of each step of the workflow are represented by dark blue rectangles. Tools automatically employed at each step are represented next to the arrows in the flow.

**Extended Data Fig. 2.**
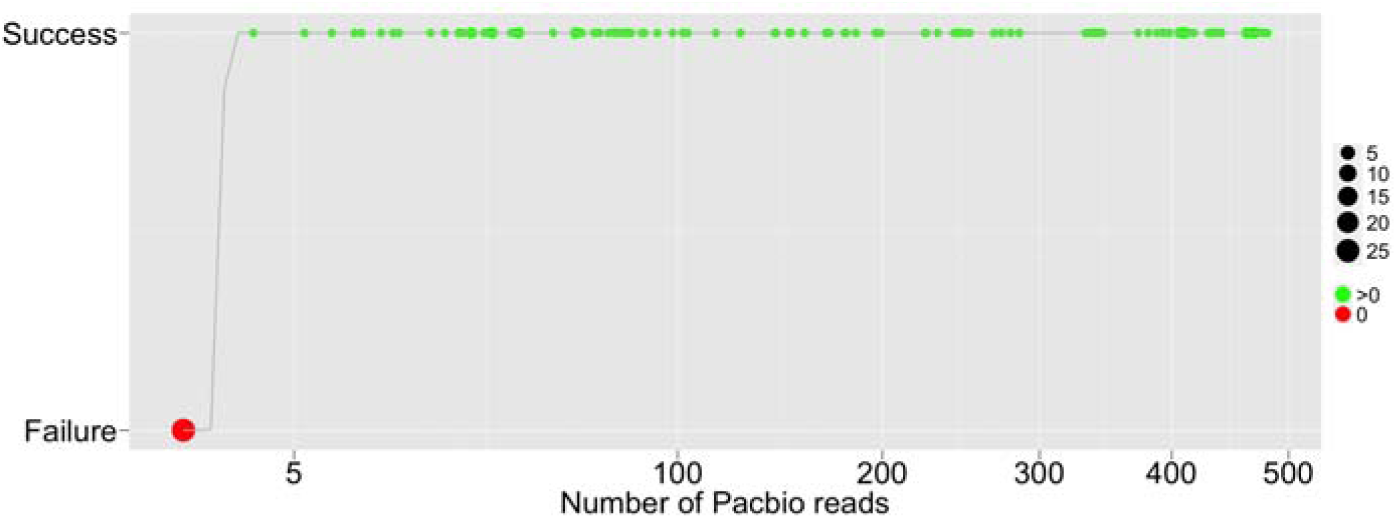
Assembly success by the availability of long mtDNA reads. The x axis is in sqrt to highlight success rate in the lower ranger. 0 read counts highlighted in red, >0 in green. The fitted curve corresponds to a generalized linear model Success ∼ Number of long mtDNA reads using a binomial distribution.

**Extended Data Fig. 3.**
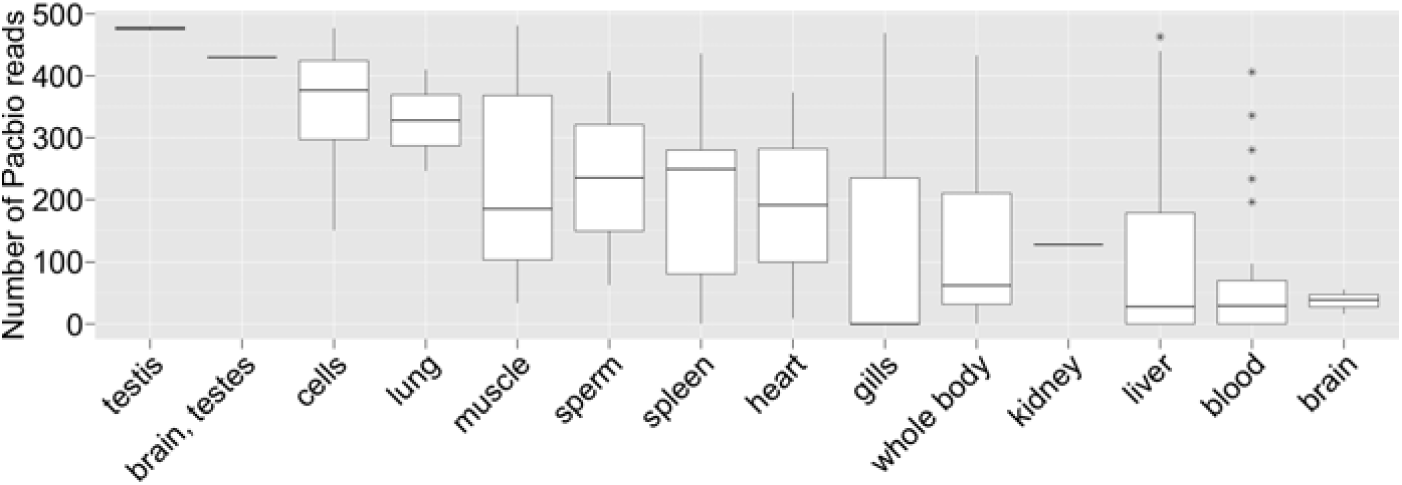
PacBio CLR mitochondrial read counts in different tissues. Black bar, average. Hinges correspond to the first and third quartiles. Whisker extends from the hinge to the values no further than 1.5 x IQR from the hinges. N = 100. *, outliers.

**Extended Data Fig. 4.**
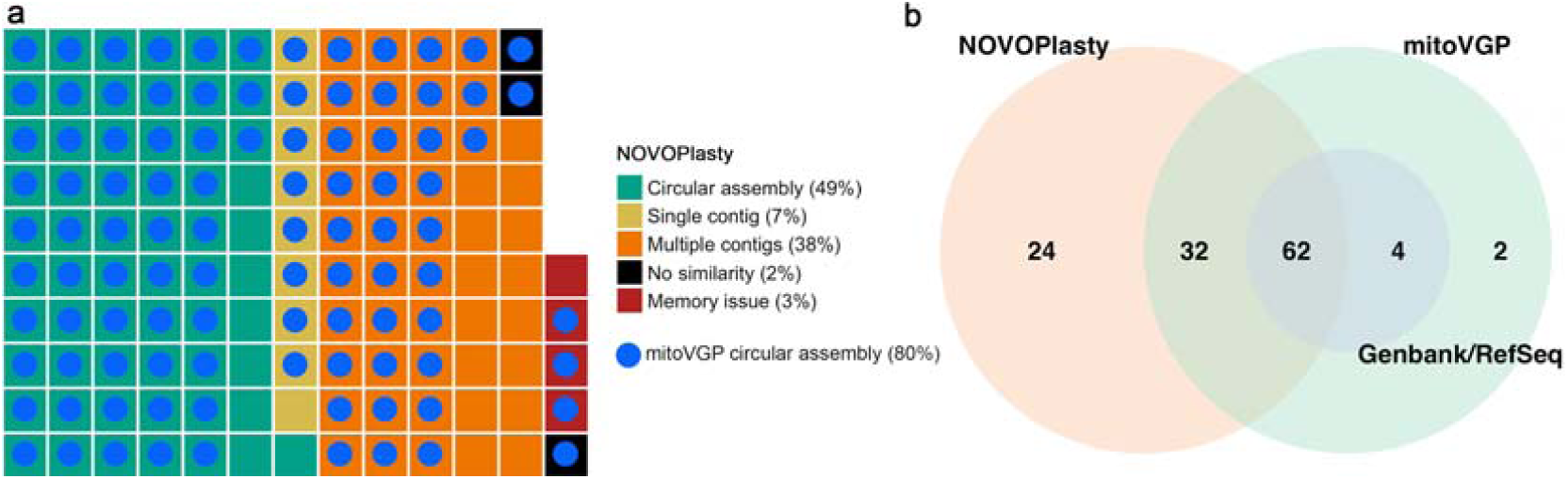
mitoVGP assembly results and comparisons. **a**, Benchmarking of the mitoVGP results with the short-read organelle genome assembler NOVOplasty. Of the successful assemblies, 49% of NOVOPlasty assembled as circular according to the software, 7% in a single contig (yellow), and 38% assembled in multiple contigs (orange). Three assemblies showed little or no similarity with corresponding reference and mitoVGP assemblies and were excluded from the analyses (black). The assembly failed in four species due to reproducible software errors (red). Matching successful mitoVGP assemblies are highlighted by the blue circles. **b**, Venn diagram of successful assemblies in the three datasets.

**Extended Data Fig. 5.**
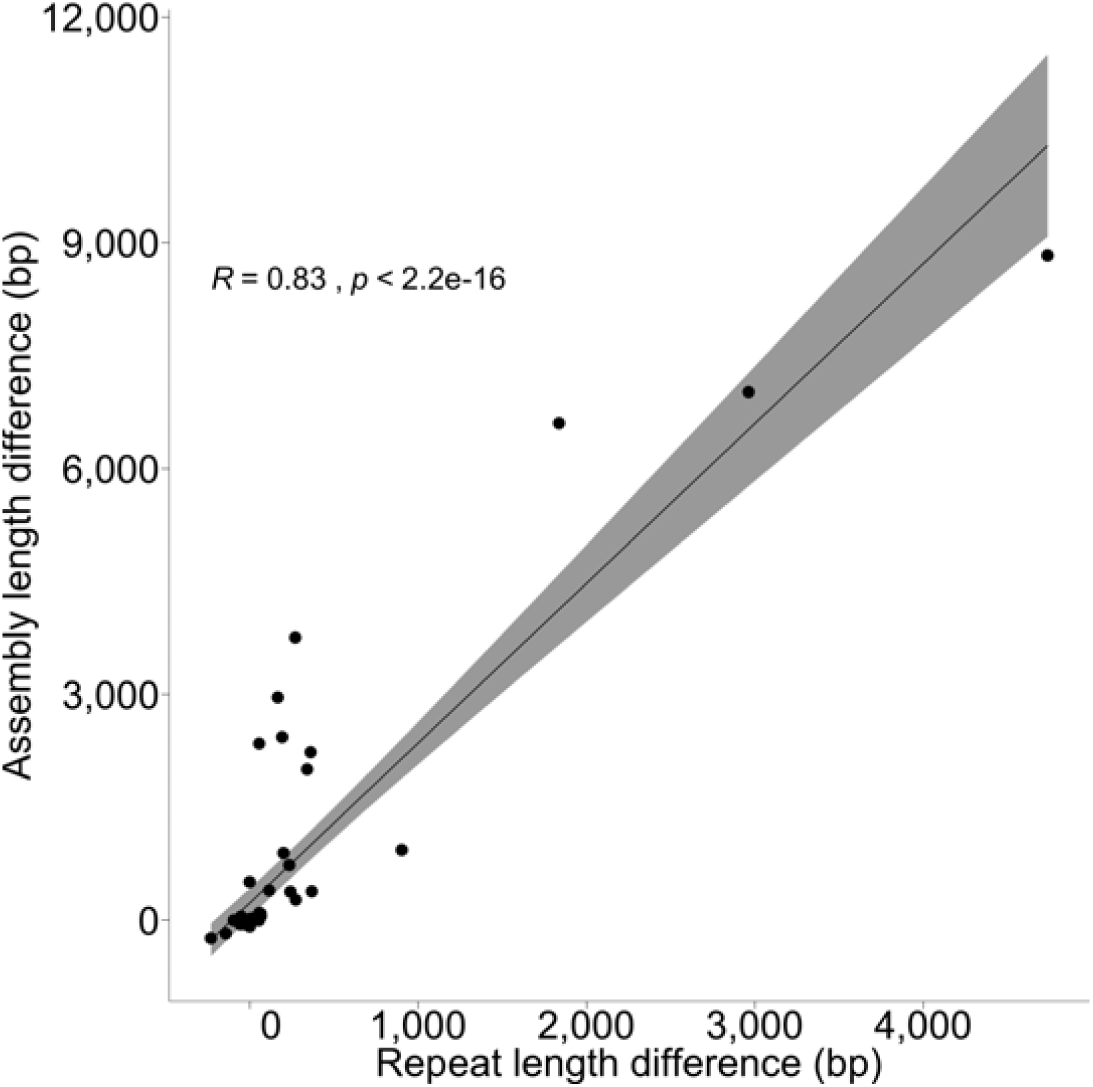
Correlation between differences in repeat content and assembly length between the mitoVGP versus the Genbank/Refseq assemblies. Regression line and confidence intervals are shown. Statistics: Spearman ρ correlation.

**Extended Data Fig. 6.**
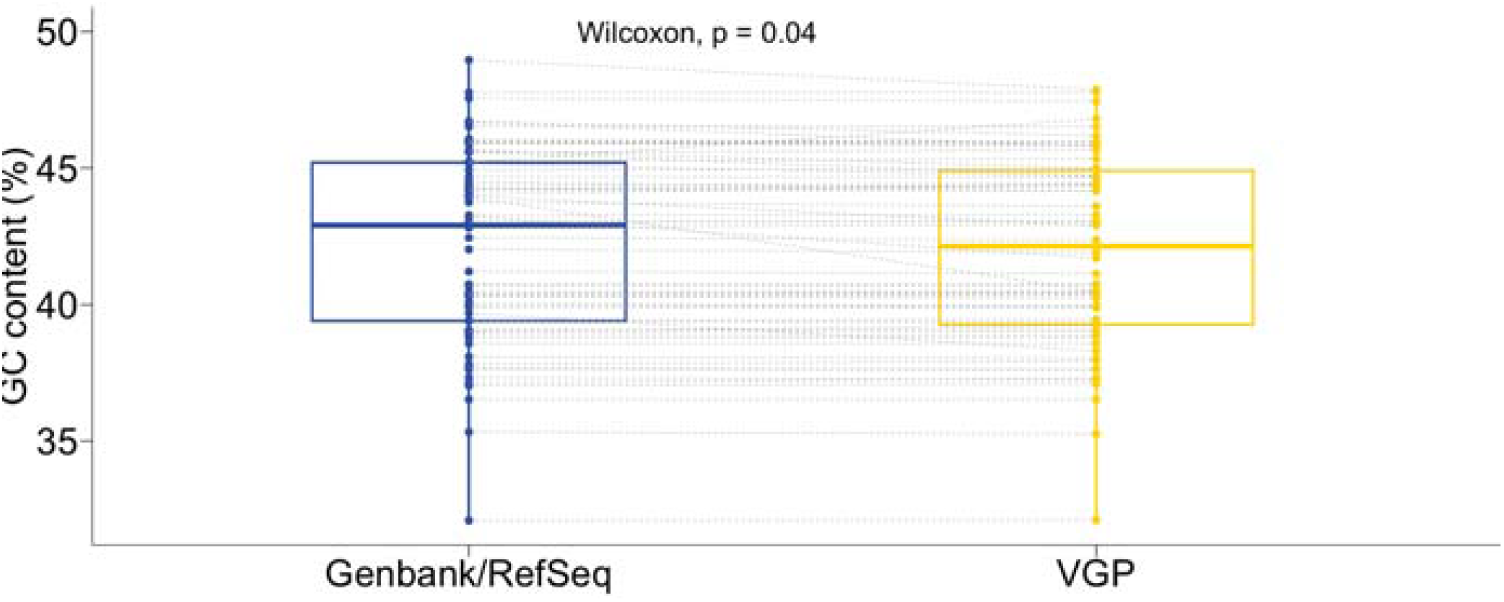
Paired comparisons of GC content between the mitoVGP assemblies and their Genbank/RefSeq counterparts. On average, the GC content was slightly lower in the mitoVGP assembly compared to the Genbank/REfSeq counterpart. Two-sided Wilcoxon test.

**Extended Data Fig. 7.**
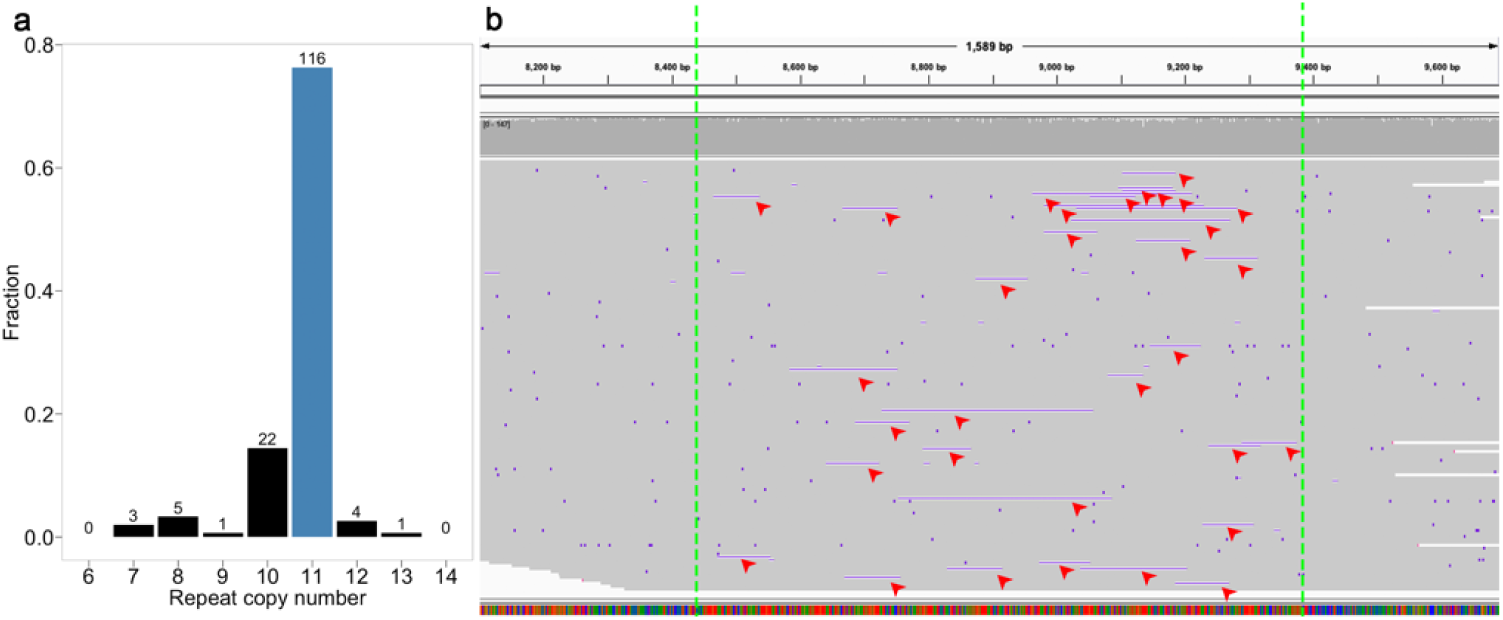
Evidence of heteroplasmy associated with a tandem repeat in the Kakapo mitochondrial genome. **a**, Fraction of CLR reads that support the copy number in the MitoVGP reference (blue bar is 11 copies). **b**, IGV^67^ plot showing the PacBio CLR alignment of reads that fully span the ∼925 bp-long tandem repeat (between green dashed lines, repeat unit = 84 bp), highlighting the presence of reads that support the copy number of 11 in the mitoVGP reference, but also reads supporting fewer copies of the repeat (red arrows, black in panel a).

**Extended Data Fig. 8.**
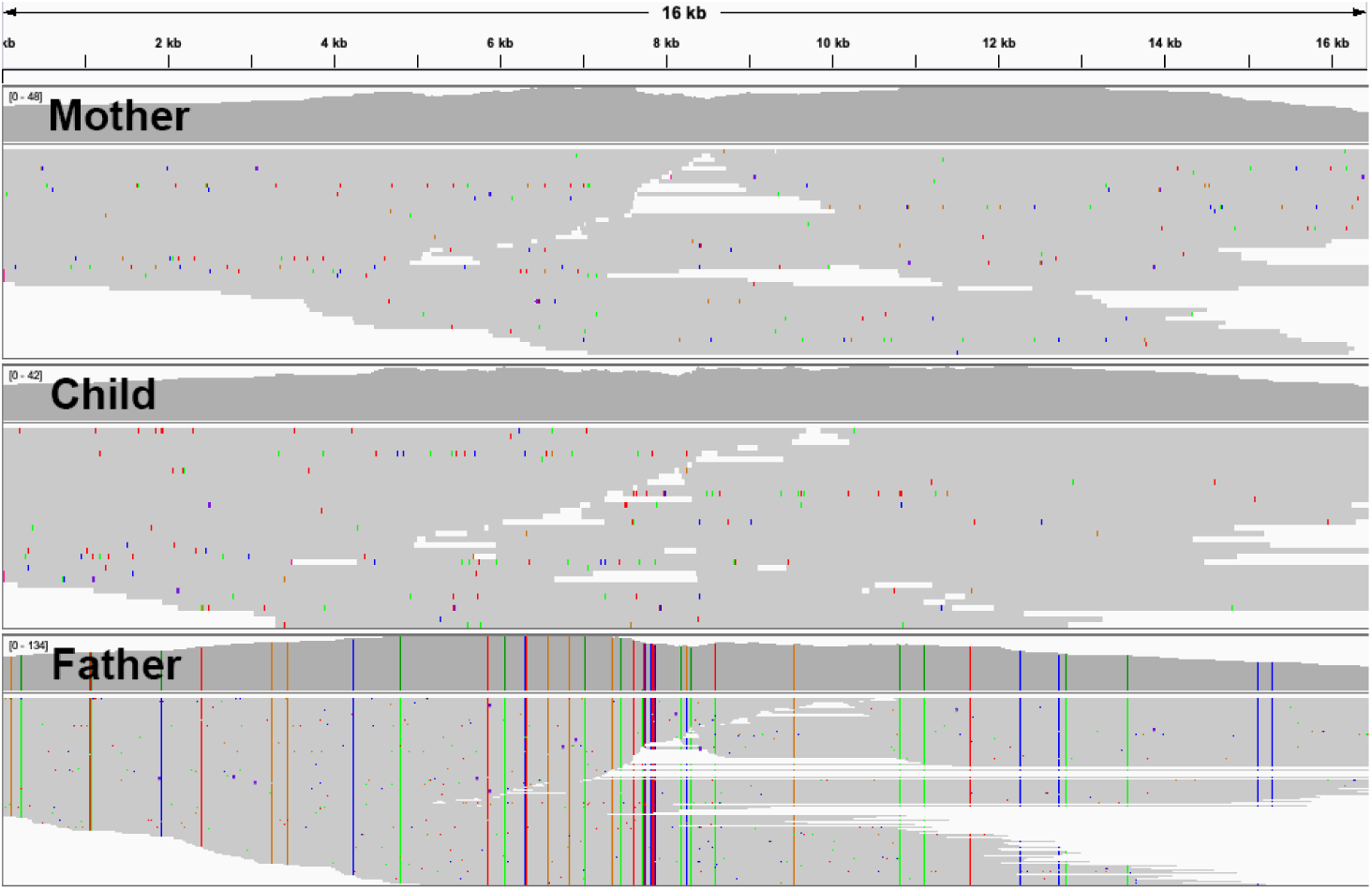
Long accurate HiFi reads from the VGP human trio mapped to the child mitogenome assembly. MtDNA CLR alignments in the mitoVGP human reference, based on trio data, showing uniform long-read coverage across the reference, highlighting inheritance patterns from the mother. First window, mother. Second window, child. Third window, father. Indels < 5 bp are masked.

**Supplementary Note 1**

Given the original ratio between mtDNA and nDNA ΔL, the relative increase in the size of the nuclear genome length as well as in the total amount of raw data to attain the same coverage ΔS, the relative gain in the number of mtDNA reads ΔR_M_ is given by the formula:

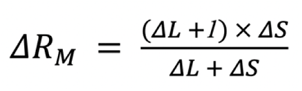

For example, assuming a typical vertebrate mitogenome size of 17 kbp, a mitochondria-rich tissue with 1,000 mitochondria per cell, when sequencing a nuclear genome three times bigger at the same coverage (e.g. a mammalian genome vs a bird genome) the expected relative gain in mtDNA reads is only 1.3%. Given that mitochondria-rich tissues would also invariably generate abundant mtDNA reads, this small difference clearly points to factors other than the total raw Gbp in the dataset, including the accuracy in estimating the nuclear genome size, predominantly driving the abundance of mtDNA reads.

